# Ultraviolet Rate Constants of Pathogenic Bacteria: A Database of Genomic Modeling Predictions

**DOI:** 10.1101/2022.05.26.493671

**Authors:** Wladyslaw Kowalski, William P. Bahnfleth, Normand Brais, Thomas J. Walsh

## Abstract

A database of bacterial ultraviolet (UV) susceptibilities is developed from an empirical model that correlates genomic parameters with UV rate constants. Software is used to count and evaluate potential ultraviolet photodimers and identifying hot spots in bacterial genomes. The method counts dimers that potentially form between adjacent bases that occur at specific genomic motifs such as TT, TC, CT, & CC. Hot spots are identified where clusters of three or more consecutive pyrimidines can enhance absorption of UV photons. The model incorporates nine genomic parameters into a single variable for each species that represents its relative dimerization potential. The bacteria model is based on a curve fit of the dimerization potential to the ultraviolet rate constant data for 92 bacteria species represented by 216 data sets from published studies. There were 4 outliers excluded from the model resulting in a 98% Confidence Interval. The curve fit resulted in a Pearson correlation coefficient of 80%. All identifiable bacteria important to human health, including zoonotic bacteria, were included in the database and predictions of ultraviolet rate constants were made based on their specific genomes. This database is provided to assist healthcare personnel and researchers in the event of outbreaks of bacteria for which the ultraviolet susceptibility is untested and where it may be hazardous to assess due to virulence. Rapid sequencing of the complete genome of any emerging pathogen will now allow its ultraviolet susceptibility to be estimated with equal rapidity. Researchers are invited to challenge these predictions.

**Importance:** This research demonstrates the feasibility of using the complete genomes of bacteria to determine their susceptibility to ultraviolet light. Ultraviolet rate constants can now be estimated in advance of any laboratory test. The genomic methods developed herein allow for the assembly of a complete database of ultraviolet susceptibilities of pathogenic bacteria without resorting to laboratory tests. This UV rate constant information can be used to size effective ultraviolet disinfection systems for any specific bacterial pathogen when it becomes a problem.

## Introduction

In spite of the widespread application of ultraviolet disinfection technologies in healthcare and other industries, less than 25% of pathogenic bacteria have been studied in laboratories to assess their UV rate constant. A complete database of bacteria UV rate constants would be useful for estimating the required UV dosage for bacteria when using UV disinfection equipment. Laboratory testing can be expensive, time-consuming, and even hazardous but the use of genomic modeling can enable rapid estimation of required UV dosage for any new or emerging pathogen once the genome is sequenced. Genomic predictions of UV rate constants can potentially be extremely accurate, more accurate than typical laboratory test results. The accuracy achievable by genomic modeling may be such that predictions could serve as a standard by which to judge the accuracy of laboratory testing, in much the same way as laws of physics are used to assess the efficiency of motors and the strength of structures. The present research is based on an empirical model that is fit to existing data and represents a step towards a fundamentals-based model of ultraviolet susceptibility that promises accuracy beyond six decimal places. The present research addresses bacteria because they have long been the largest and most persistent infection problem in healthcare. Although this view is being challenged by the current pandemic, the success of this bacteria genomic model will also lead to a model for predicting UV rate constants for viruses as well.

## Methods

The methods used to develop the model and make predictions are purely analytical and empirical and involved firstly a compilation of UV rate constant studies for bacteria and computation of the UV rate constants of k values. The measured UV rate constants are averaged when there are multiple studies and are applied in an empirical model that correlates the k values with a theoretical dimerization value, Dv, which quantifies a series of genomic parameters into a single value. The dimerization value is computed directly from the genomes for all subsequent bacteria to produce the predicted UV rate constant. The details of these computations are discussed in the following subsections.

### Rate Constant Studies

Studies on ultraviolet (254 nm) survival of bacteria on surfaces or in water were reviewed. Criteria for selection included having interpretable results that would define the first stage of exponential decay independent of any survival tail (the second stage) or shoulder in the survival curve, since the presence of a tail can confuse interpretation of the first stage rate constant. Table 1 summarizes the 92 bacteria that form the basis of the model and the UV rate constants averaged from the indicated studies. Every attempt was made to be inclusive of all available studies but invariably there were a few studies that defied interpretation or for which a genome was lacking. Shown in the columns of Table 1 are the UV rate constant k computed as the average of all studies, the Mean Aerodynamic Diameter (MAD), the D90 values, and the genome identifier or accession number. In some cases the identifier is the first in a series of shotgun sequence identifiers. The number of data sets reflects the number of studies up to a maximum of 3 beyond which no further weight is given to the species in the model. This avoids bias for the small number of species that have a large number of studies and 3 is an appropriate number of studies from which to obtain an average value. In cases where an excessive number of studies exist many were omitted due to redundancy. Four species formed outliers in this model and were excluded. *Nocardia asteroides* was also a low outlier and it is a bacteria that more resembles a fungus and may not belong to this data set, but more studies on *Nocardia* species are needed to resolve this issue. *Enterobacter cloacae* was a low outlier. *Mycobacterium abscessus, Burkholderia mallei*, and *B. pseudomallei* were all high outliers. These outliers were each single studies except for *B. pseudomallei* which had a second study which was not an outlier and which is included in the model in Table 1. *Micrococcus radiodurans* was expected to be a low outlier due to its unique ability to resist radiation and was excluded from consideration altogether except that it is shown for reference.

**Table 1:** Average Rate Constant Summary from Studies.

| Bacteria | Source | Data Sets | Mean $k_1$<br>$m^2/J$ | Avg $D_{90}$<br>$J/m^2$ | Mean Dia.<br>$\mu m$ | Genome ACC.#<br>NCBI, Kegg | Genome<br>kb |
| --- | --- | --- | --- | --- | --- | --- | --- |
| *Acinetobacter baumannii* | 84,97 | 2 | 0.1527 | 15 | 1.31 | NC_008801 | 3904 |
| *Aeromonas punctata (caviae)* | 56 | 1 | 0.1727 | 13 | 0.70 | NZ_AP022254 | 4566 |
| *Aeromonas hydrophila* | 56101 | 2 | 0.1528 | 15 | 0.70 | NC_008570 | 4744 |
| *Aeromonas salmonicida* | 55,66,88 | 3 | 0.1501 | 15 | 0.70 | NC_009348 | 5041 |
| *Bacillus anthracis* | 46,57 | 2 | 0.1050 | 22 | 1.41 | NC_007530 | 5504 |
| *Bacillus cereus* | 23,30 | 2 | 0.0582 | 40 | 1.28 | NC_004722 | 5427 |
| *Bacillus megaterium* | 43 | 1 | 0.0540 | 43 | 1.79 | NZ_CP009920 | 5747 |
| *Bacillus subtilis* | 40,68,69,85,100 | 3 | 0.0549 | 42 | 0.56 | NC_000964 | 4215 |
| *Bacteroides fragilis* | 51,94,89 | 3 | 0.1020 | 23 | 1.15 | NC_003228 | 5242 |
| *Brevundimonas diminuta* | 1 | 1 | 0.0311 | 74 | 0.27 | NZ_UIGE01000002 | 3439 |
| *Brucella melitensis* | 87 | 1 | 0.0722 | 32 | 0.64 | NZ_WUEE01000001 | 3291 |
| *Brucella suis* | 87 | 1 | 0.1057 | 22 | 0.64 | NZ_CP007719 | 3297 |
| *Burkholderia cenocepacia* | 1 | 1 | 0.0397 | 58 | 0.76 | NZ_MUTR01000001 | 9961 |
| *Burkholderia cepacia* | 1,15 | 2 | 0.0508 | 45 | 0.76 | NZ_CP045235 | 8367 |
| *Burkholderia pseudomallei* | 49 | 1 | 0.0461 | 50 | 0.84 | NC_006350 | 7248 |
| *Campylobacter jejuni* | 14,101 | 2 | 0.1375 | 17 | 0.63 | NC_002163 | 1641 |
| *Citrobacter freundii* | 35,106 | 2 | 0.0525 | 44 | 1.28 | NZ_AOMS01000001 | 4900 |
| *Citrobacter koseri* | 35 | 1 | 0.0461 | 50 | 1.28 | NC_009792 | 4735 |
| *Clostridium perfringens* | 44 | 1 | 0.0600 | 38 | 5.00 | NC_003366 | 3031 |
| *Corynebacterium diphtheria* | 92 | 1 | 0.0680 | 34 | 1.06 | NZ_LN831026 | 2464 |
| *Coxiella burnetii* | 67,69 | 2 | 0.1036 | 22 | 0.32 | NC_002971 | 2033 |
| *Enterobacter cloacae* | 106 | 1 | 0.0461 | 50 | 1.41 | NZ_CP008823 | 5503 |
| *Enterococcus faecalis* | 1,19,20,38,40,58,77 | 3 | 0.0533 | 43 | 1.11 | NZ_KE351595 | 2929 |
| *Enterococcus faecium* | 72,74,75 | 3 | 0.0639 | 36 | 1.11 | NC_017958 | 3053 |
| *Escherichia coli* | 12,24,27,31,83,92,99,102 | 3 | 0.0973 | 24 | 0.94 | NC_004431 | 5231 |
| *Fluoribacter bozemaniae* | 36,57,103 | 3 | 0.1587 | 15 | 0.73 | NZ_UGGY01000007 | 4232 |
| *Fluoribacter dumoffii* | 36,57 | 2 | 0.1039 | 22 | 0.73 | NZ_LNXZ01000001 | 3817 |
| *Fluoribacter gormanii* | 36,57 | 2 | 0.1337 | 17 | 0.73 | NZ_UGGV01000002 | 3873 |
| *Francisella tularensis* | 87 | 1 | 0.1708 | 13 | 0.22 | NZ_KN046815 | 1818 |
| *Haemophilus influenzae* | 6,76 | 3 | 0.1265 | 18 | 0.33 | NC_000907 | 1830 |
| *Helicobacter pylori* | 40 | 1 | 0.0971 | 24 | 0.94 | NC_000921 | 1644 |
| *Klebsiella pneumoniae* | 35,106 | 2 | 0.0504 | 46 | 0.55 | NC_009648 | 5315 |
| *Lactococcus lactis* | 85 | 1 | 0.0557 | 41 | 0.85 | NZ_LT599049 | 2731 |
| *Legionella jordanis* | 57 | 1 | 0.0768 | 30 | 0.73 | NZ_LR134383 | 3134 |
| *Legionella longbeachae* | 18,36 | 2 | 0.2022 | 11 | 0.73 | NC_013861 | 4149 |
| *Legionella oakridgensis* | 57 | 1 | 0.0677 | 34 | 0.73 | NZ_CP004006 | 2773 |
| *Legionella pneumophila* | 3,18,36,57,79,101,103 | 3 | 0.1447 | 16 | 0.73 | NC_006369 | 3346 |
| *Legionella wadsworthii* | 57 | 1 | 0.1212 | 19 | 0.73 | NZ_UGPB01000002 | 3603 |
| *Listeria monocytogenes* | 24,55,104 | 3 | 0.0771 | 30 | 0.71 | NC_002973 | 4844 |
| *Lysinibacillus fusiformis* | 85 | 1 | 0.0763 | 30 | 0.56 | NZ_CP010820 | 2905 |
| *Micrococcus luteus* | 4 | 1 | 0.0237 | 97 | 0.39 | NC_012803 | 2501 |
| *Moraxella catarrhalis* | 85 | 1 | 0.0768 | 30 | 0.77 | NC_014147 | 1863 |
| *Mycobacterium avium* | 41,73,93 | 3 | 0.0429 | 54 | 0.76 | NC_008595 | 5475 |
| *Mycobacterium bovis BCG* | 24 | 1 | 0.1007 | 23 | 0.55 | NZ_CP039850 | 4412 |
| *Mycobacterium gilvum* | 27 | 1 | 0.0285 | 81 | 0.78 | NC_008595 | 5620 |
| *Mycobacterium intracellulare* | 27,41 | 2 | 0.0293 | 79 | 0.76 | NC_016946 | 5402 |
| *Mycobacterium kansasii* | 27 | 1 | 0.0408 | 56 | 0.54 | NC_022663 | 6577 |
| *Mycobacterium marinum* | 27 | 1 | 0.0468 | 49 | 0.55 | NC_010612 | 6637 |
| *Mycobacterium smegmatis* | 11,27 | 3 | 0.0359 | 64 | 0.60 | NC_008596 | 6988 |
| *Mycobacterium terrae* | 10 | 1 | 0.0461 | 50 | 0.64 | CP002329 | 4644 |
| *Mycobacterium tuberculosis* | 11,24,25,27 | 3 | 0.0747 | 31 | 0.55 | NC_009565 | 4424 |
| *Mycolicibacterium fortuitum* | 27 | 2 | 0.0542 | 42 | 0.80 | NZ_CP011269 | 6255 |
| *Mycolicibacterium phlei* | 24,27 | 2 | 0.0549 | 42 | 0.50 | NZ_CP014475 | 5350 |

**Table 1: Average Rate Constant Summary from Studies**
| Bacteria | Source | Data Sets | Mean $k_1$<br>$m^2/J$ | Avg $D_{90}$<br>$J/m^2$ | Mean Dia.<br>$\mu m$ | Genome ACC.#<br>NCBI, Kegg | Genome<br>kb |
| --- | --- | --- | --- | --- | --- | --- | --- |
| Mycoplasma arthritidis | 32 | 1 | 0.1050 | 22 | 0.24 | NC_011025 | 820 |
| Mycoplasma fermentans | 32 | 1 | 0.1163 | 20 | 0.31 | NC_014921 | 1119 |
| Mycoplasma hominis Type 1 | 32 | 1 | 0.1396 | 17 | 0.19 | NC_013511 | 695 |
| Mycoplasma orale Type 1 | 32 | 1 | 0.1308 | 18 | 0.24 | NZ_LR214940 | 806 |
| Mycoplasma pneumoniae | 32 | 1 | 0.1396 | 17 | 0.31 | NZ_CP010546 | 817 |
| Mycoplasma salivarium | 32 | 1 | 0.1308 | 18 | 0.32 | NZ_LR214938 | 1737 |
| Neisseria gonorrhoeae | 16 | 1 | 0.0695 | 33 | 0.77 | NZ_LT591897 | 2292 |
| Proteus mirabilis | 44 | 1 | 0.2890 | 8 | 0.63 | NC_008806 | 4064 |
| Proteus vulgaris | 85 | 1 | 0.1085 | 21 | 0.63 | NZ_LR590468 | 3925 |
| Pseudomonas aeruginosa | 1,9,13,31,35,39,74 | 3 | 0.0751 | 31 | 0.88 | NC_002516 | 6588 |
| Pseudomonas fluorescens | 4,55,85 | 3 | 0.0919 | 25 | 0.69 | NC_007492 | 6846 |
| Raoultella terrigena | 101 | 1 | 0.0700 | 33 | 0.70 | NZ_LANE01000008 | 5717 |
| Rickettsia prowazekii | 2 | 1 | 0.1760 | 13 | 0.60 | NC_008807 | 1112 |
| Salmonella enterica subsp anatum | 98 | 1 | 0.0384 | 60 | 1.12 | NC_003197 | 4831 |
| Salmonella enterica subsp derby | 98 | 1 | 0.0636 | 36 | 1.12 | CP029486 | 4885 |
| Salmonella enteritidis | 24,57,98 | 3 | 0.0671 | 34 | 1.12 | CP032851 | 4680 |
| Salmonella infantis | 98 | 1 | 0.1151 | 20 | 1.12 | CP038507 | 4728 |
| Salmonella typhi | 19,92,101 | 3 | 0.1122 | 21 | 1.12 | NC_003198 | 5134 |
| Salmonella typhimurium | 19,48,54,62,98 | 3 | 0.1002 | 23 | 1.12 | NC_003197 | 4951 |
| Serratia marcescens | 7,24,38,39,85,92,105 | 3 | 0.1111 | 21 | 0.72 | NZ_HG326223 | 5114 |
| Shewanella algae | 82 | 1 | 0.2709 | 8 | 0.51 | NZ_LN810009 | 5003 |
| Shewanella putrefaciens | 1,82 | 2 | 0.3587 | 6 | 0.51 | NC_009438 | 4659 |
| Shigella dysenteriae | 48,92,101 | 3 | 0.1236 | 19 | 0.63 | NC_009344 | 4369 |
| Shigella sonnei | 19 | 1 | 0.1486 | 15 | 0.63 | NC_007384 | 4825 |
| Staphylococcus aureus | 1,19,23,28,33,74,92 | 3 | 0.0929 | 25 | 0.77 | NC_010079 | 2873 |
| Staphylococcus epidermis | 24,92 | 2 | 0.0875 | 26 | 0.87 | NC_004461 | 2617 |
| Stenotrophomonas maltophilia | 1,15 | 2 | 0.0447 | 52 | 0.60 | NC_010943 | 4851 |
| Streptococcus pyogenes | 92 | 1 | 0.1084 | 21 | 1.02 | NC_002737 | 1915 |
| Streptococcus viridans | 92 | 1 | 0.1134 | 20 | 1.00 | NZ_UHHS01000002 | 1997 |
| Streptomyces coelicolor | 49 | 1 | 0.0384 | 60 | 1.00 | NC_003888 | 8668 |
| Streptomyces griseus | 49,52 | 2 | 0.0281 | 82 | 0.71 | NC_008802 | 8546 |
| Tatlockia micdadei | 35,56 | 1 | 0.1070 | 22 | 0.73 | NZ_LN614830 | 3310 |
| Vibrio anguillarum | 56,65,88 | 3 | 0.1644 | 14 | 0.60 | NC_015633 | 4052 |
| Vibrio cholerae | 5,101 | 3 | 0.1957 | 12 | 0.76 | NZ_AP014524 | 4031 |
| Vibrio ordalii | 88 | 1 | 0.1256 | 18 | 0.60 | NZ_AJYS02000001 | 3895 |
| Vibrio parahaemolyticus | 5,78 | 2 | 0.1797 | 13 | 0.77 | NZ_CP031781 | 3289 |
| Yersinia enterocolitica | 14,127,23,101 | 3 | 0.1482 | 16 | 0.76 | NC_008800 | 4616 |
| Yersinia pestis | 87 | 2 | 0.1645 | 14 | 0.73 | NC_003143 | 4830 |
| Yersinia ruckeri | 65 | 1 | 0.1175 | 20 | 0.87 | NZ_CP011078 | 3700 |
| Burkholderia mallei | 87 | 1 | 0.2111 | 11 | 0.87 | NC_006348 | 5836 |
| Burkholderia pseudomallei | 87 | 1 | 0.1829 | 13 | 0.84 | NC_006350 | 7248 |
| Mycobacteroides abscessus | 96 | 1 | 0.1727 | 13 | 0.55 | NZ_AP018436 | 5080 |
| Nocardia asteroides | 22 | 1 | 0.0082 | 280 | 1.41 | NZ_LR134352 | 6955 |
| Deinococcus radiodurans | 1,4,91 | 3 | 0.0039 | 593 | 1.41 | NZ_CP031500 | 3283 |

The Average UV Rate Constant k shown in Table 1 is computed from information provided in the referenced studies. If only a D90 value is provided and there is no significant shoulder or tail evident then the value is converted to a rate constant according to equation (1).

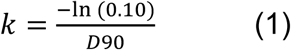

In most cases the original data is inspected to determine the UV rate constant separate from any shoulder or tail. The range of data points within which the value of k1 can be accurately computed is illustrated in Figure 2. Any points above and beyond the shoulder region, if any, and any points beyond the survival tail, if any, are suitable for use in computing the UV rate constant. In all case there are multiple studies available, the rate constant is computed as the average and not the D90 value.

**Figure 1:**
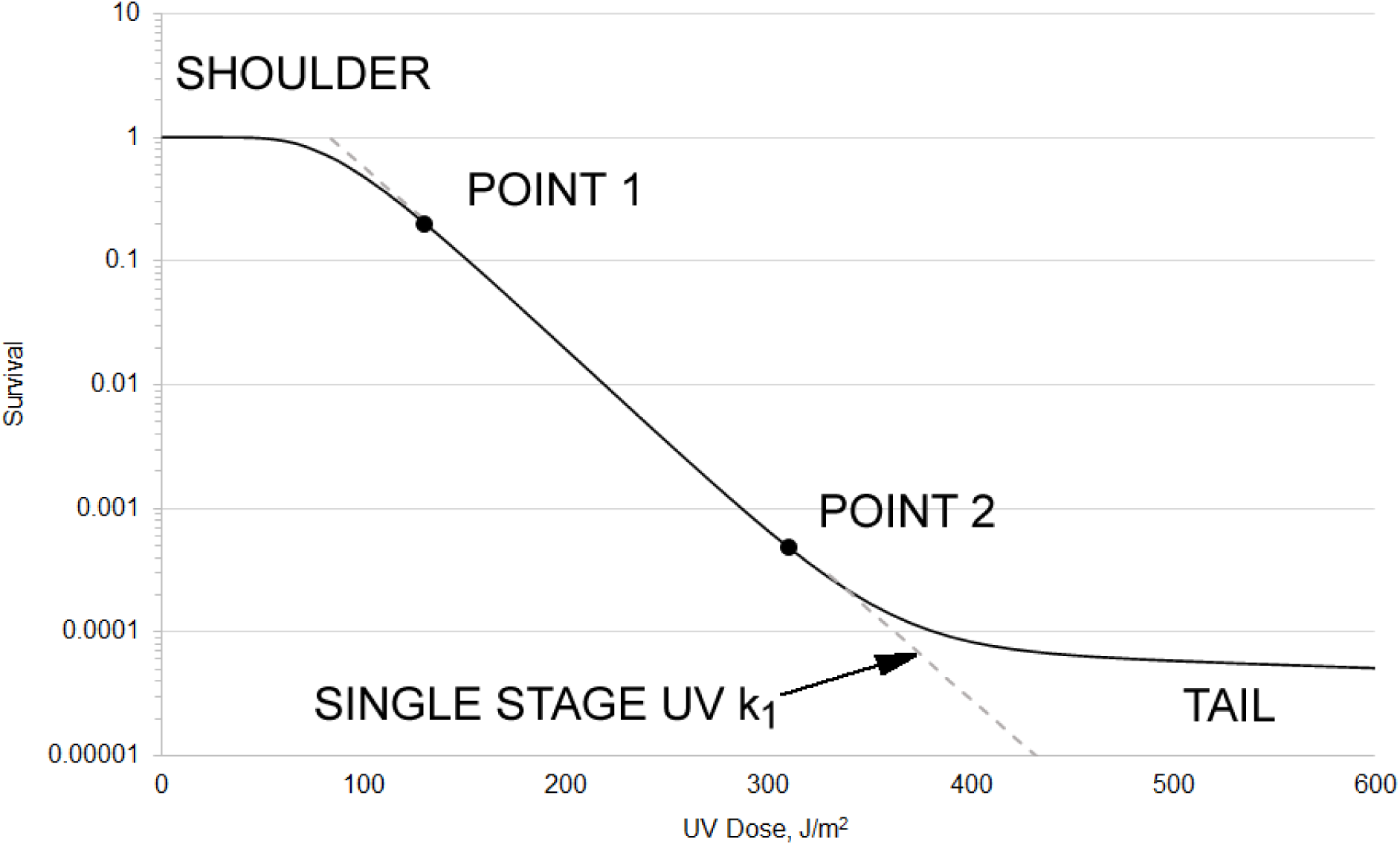
Figurative survival curve with a shoulder and a tail. The Single Stage UV rate constant k_1_ must be computed between the data points beyond the shoulder region (Point 1) and the data points before the tail develops (Point 2). The UV Dose scale shown is arbitrary.

**Figure 2:**
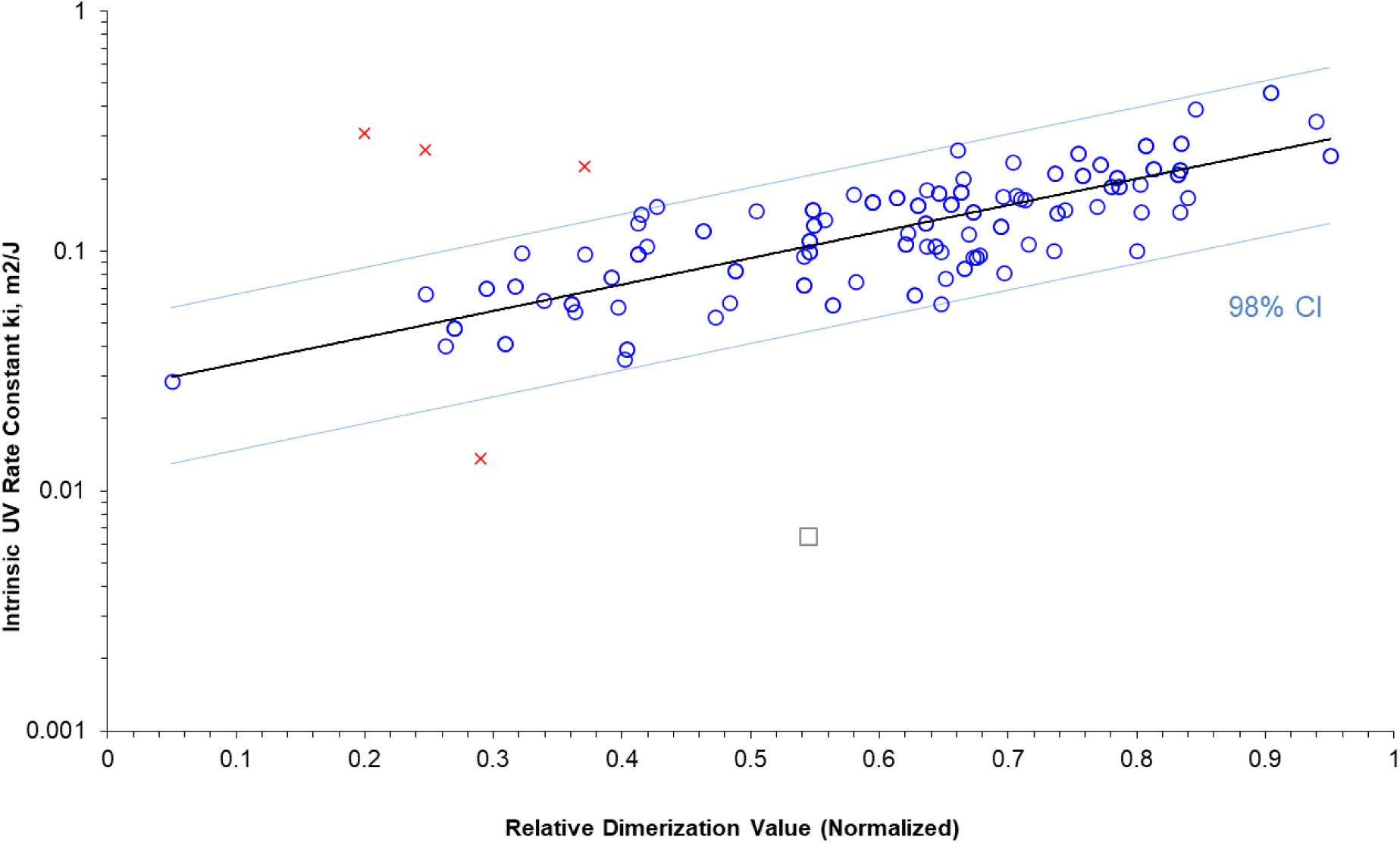
Plot of the 92 tested bacteria (blue circles) and the genomic model (dark line) with the 98% Confidence Interval (CI) high and low limits shown in light blue. Four outliers are shown as red X’s. *D. radiodurans* (hollow square) is not a pathogen but is shown for perspective since it is the most radiation resistant microbe.

The Mean Diameter listed in Table 1 represents the logmean diameter of spherical cells and for rods or oblong cells it represents the logmean diameter computed with the Cauchy Theorem, which represents the average area of the profile of the rod. The profile area accounts for the area upon which the UV rays impinge and this determines the amount of UV scattering (Kowalski 2020). The mean diameter is used to compute the shielding constant Sc, which represents the amount of shielding a cell has from UV due to its physical size. The shielding constant is used in turn to compute the intrinsic UV rate constant which is the value correlated by the empirical model. The advantage of using the intrinsic rate constant, however theoretical it may be, is seen in the fact that the MAD exponent in this model converged to zero and ceased to have any predictive value. This simplification separates the only physical parameter (MAD) from the genomic parameters and renders the core model purely genomic in nature.

Calculations of the UV rate constants from the studies in Table 1 will be made available as supplemental files online at XXXX.later.

The last five microbes shown in gray cells are the four outliers and *D. radiodurans*.

#### The Genomic Parameters

Theoretically, the exact sequence of adjacent base pairs are the primary determinants of the intrinsic UV sensitivity and the frequency of dimer formation is in the order TT>TC>CT>CC (Setlow 1966, Becker 1989, Khoe 2018). Specifically, adjacent pyrimidines have the potential to form dimers, and some purines flanked by pairs of pyrimidines may also form dimers (Pan 2011). Pyrimidine doublets, or pairs of pyrimidines, may have dimer formation suppressed when they are flanked by purines (Law 2013). Table 2 summarizes the genomic motifs that are likely to dimerize and those that are not.

**Table 2:**
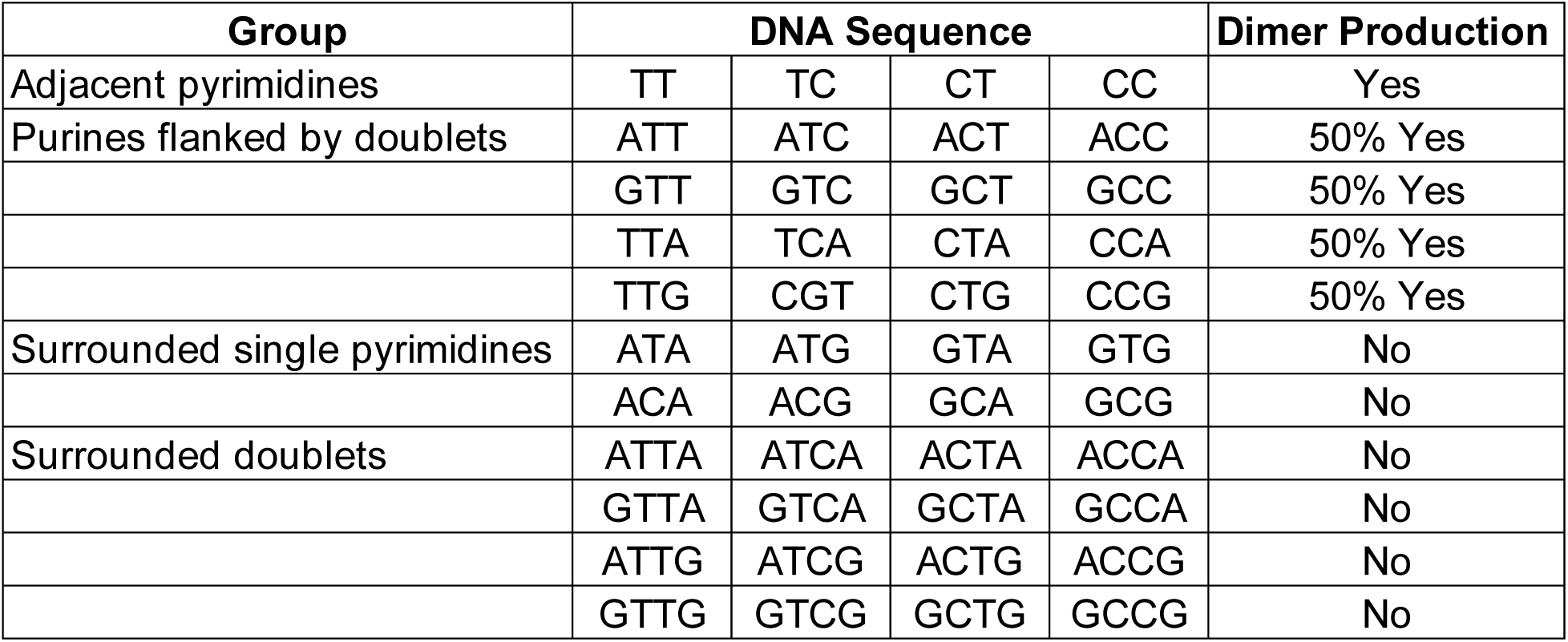
Potential Dimerization Sequences.

There are five groups of potential dimers, TT, TC, CT, CC, and YR (where R represents purines and Y represents pyrimidines). The sites are counted inclusively, wherein, for example, the sequence TT counts as 2 dimer sites and the sequence CTC counts as 3 dimer sites. Where sequences overlap, dimers are categorized based on the priority TT>TC>CT>CC>YR.

When multiple pyrimidines occur sequentially they may create a hot spot where the likelihood of dimerization is increased by a factor of up to 2 (Becker 1989). This phenomenon is known as hyperchromicity. Not enough is known about the hyperchromic effect to quantify it precisely but a parameter is included to add to the sum of dimers of any doublet or triplet whenever 3 or more pyrimidines are found in sequence (Kowalski et al 2009). This factor, H, increases the probability of dimerization for doublets and triplets based on how many adjacent pyrimidines are present in the genome, up to a value of 8 in a row where the effect levels off. The value of H may typically double the value of an isolated doublet, based on studies by Becker and Wang (1989).

The value of TTh represents the additional effect from a local hotspot, a value from 0 – 1.0 which represents the increase in dimerization probability due to being part of a cluster. The increased photoreactivity is based on an assumed sigmoid function relating the hyperchromicity factors (TTh, etc.) to the number of pyrimidines in sequence, as shown in Table 3.

**Table 3:**
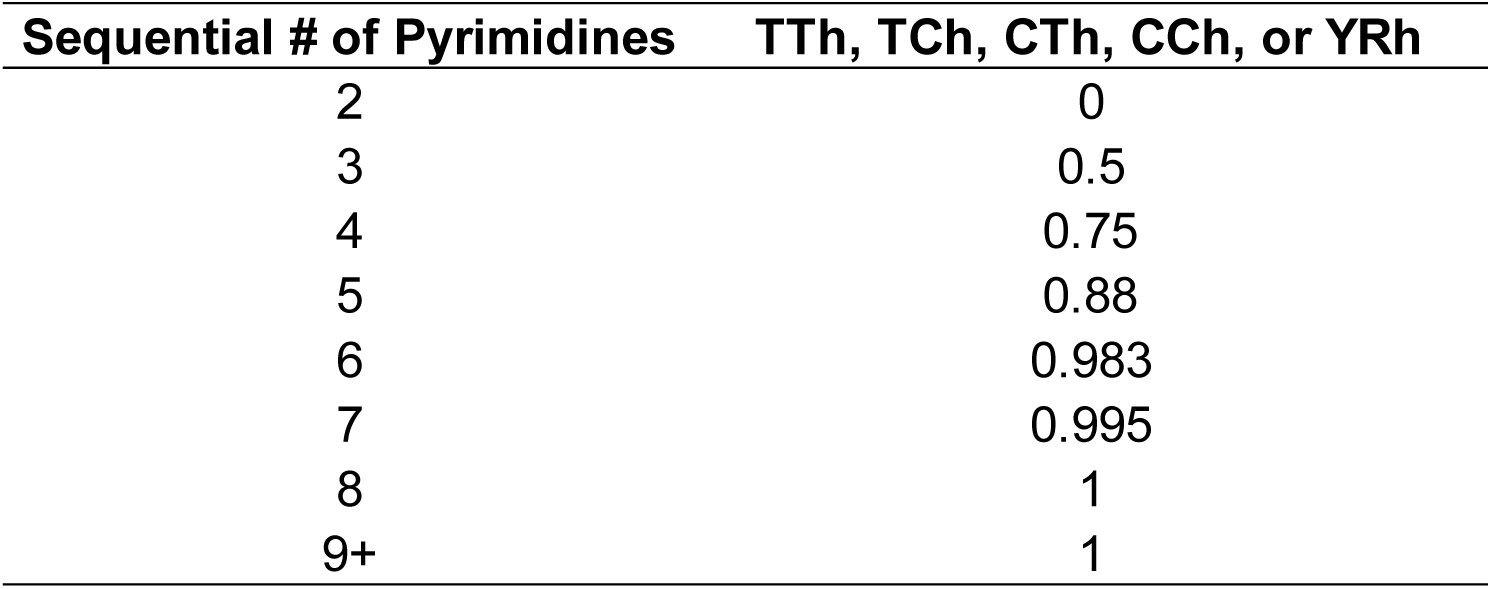
Hyperprimer Values.

It was found in the course of this research that the hyperprimers (labeled TTh, TCh, CTh, CCh & YRh) were better predictors than the specific sequences of base pairs (TT, TC, CT, CC & YR). This innovation marks a departure from previous genomic models by the author (Kowalski 2009, 2009a, 2009b, 2009c). Since the hyperprimer values represent hot spots where dimerization is amplified, this approach is called a Hot Spot model. The values of the hyperprimers were simply substituted in place of the dimer counts in the equation for computing the Dimerization value.

#### The Dimerization Value

The nine genomic parameters that proved most significant are identified in Table 3. The Relative Dimerization Value Dv is computed by multiplying all parameters together, with each having an adjustable exponent, as follows:

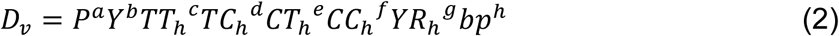

where a thru h are adjustable exponents

The exponents were adjusted through multiple iterations to maximize the least squares curve fit of ki to Dv. If the exponent of a parameter is zero, then the dimer contribution becomes unity, and the factor makes no contribution to the model. All parameters except bp are fractions since they are individually normalized as a fraction of the total bp. Since all the variables in equation (2) are dimensionless except bp, the dimerization value Dv has the units of bp.

Table 4 summarizes the final iterated values of the genomic parameter exponents after curve-fitting to maximize the Pearson Correlation coefficient.

**Table 4:**
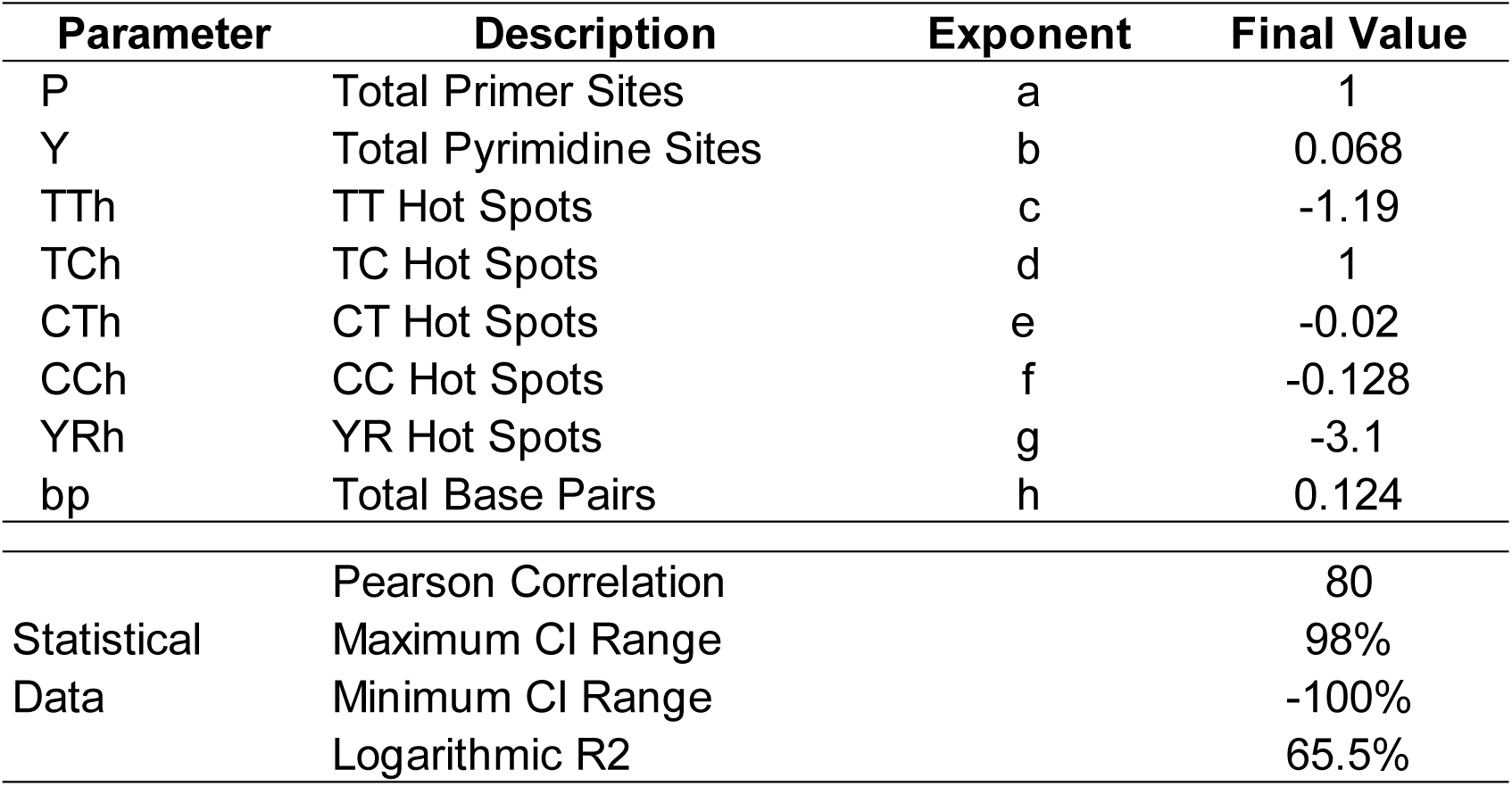
Genomic Constants of the Model and Correlations.

In this model, as opposed to previous models by the author, the intrinsic UV rate constant is used as the predictor (Kowalski 2020). The intrinsic UV rate constant is theorized as a parameter that depends solely on the genomic sequences and not on any physical parameter. The physical size of a microorganism plays a role in determining its ultraviolet sensitivity via Mie scattering effects. The intrinsic UV rate constant is defined as:

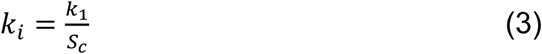

Where k_1_ = first stage UV rate constant, m^2^/J

S_c_ = shielding constant

The shielding constant is computed as the complement of the ratio of the scattering cross-section to the extinction cross-section per Kowalski (2020):

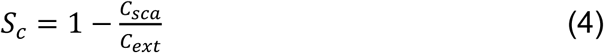

Computations of the shielding constants are provided in the supplementary materials. UV rate constants from Table 1 are converted to intrinsic rate constants for use in the genomic model. The resulting predictions are then converted back to first stage rate constants in the final summary.

#### The Empirical Model

The final empirical model correlating measured UV rate constants with those predicted by the model is summarized by the parameters in Table 3 and plotted in Figure 2.

The complete equation for the intrinsic rate constant ki can be written as follows:

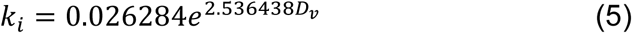

Equations (2) and (3) can then be combined into a complete closed form equation to determine k_i_.

#### The Bacteria Database

Table 5 presents a summary of all human pathogens and zoonotic pathogens that could be identified that have a published genome. The Type of microbe (H=Human, Z = Zoonotic) is specified for each pathogen. The “Genome RefSeq/NCBI#” shows either the complete genome number or in some cases it shows the first chromosome or segment identification number from a shotgun sequence. The UV k is given along with the D90 range (for a 98% Confidence Interval) and the Mean D90 is computed based on the UV k. No account is taken for shoulders in this summary and therefore the UV k value and the D90 value represent the absence of any shoulder.

**Table 5:**
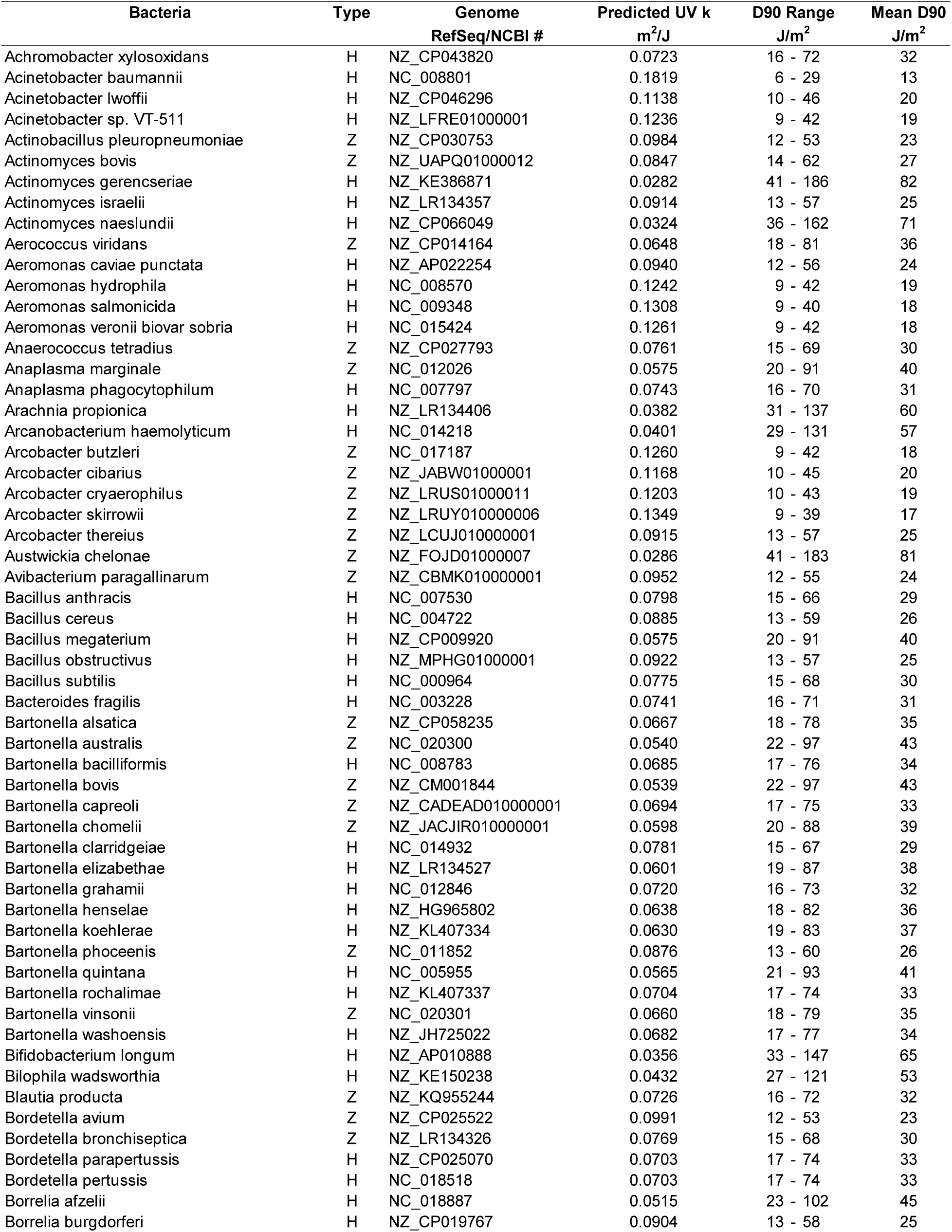
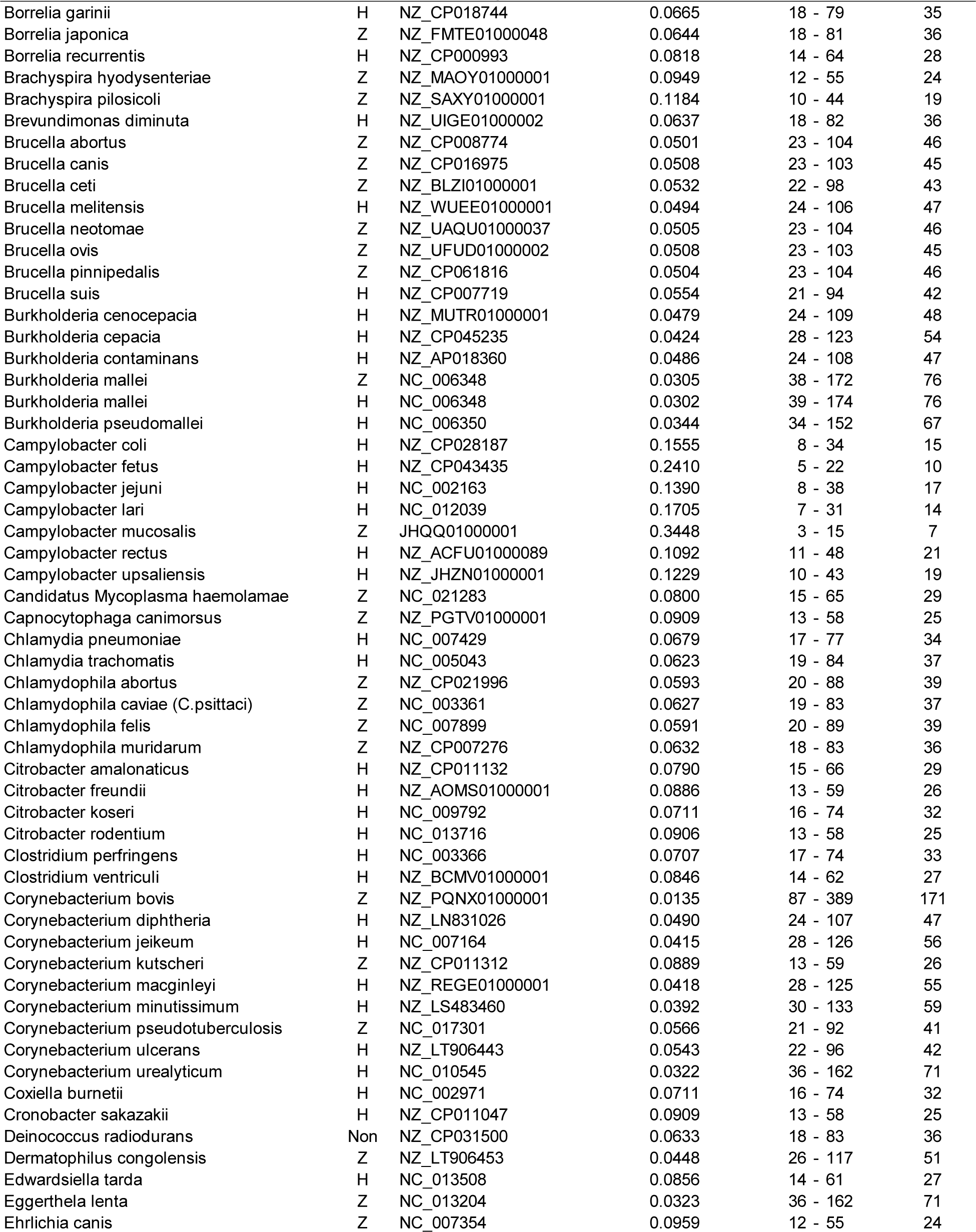
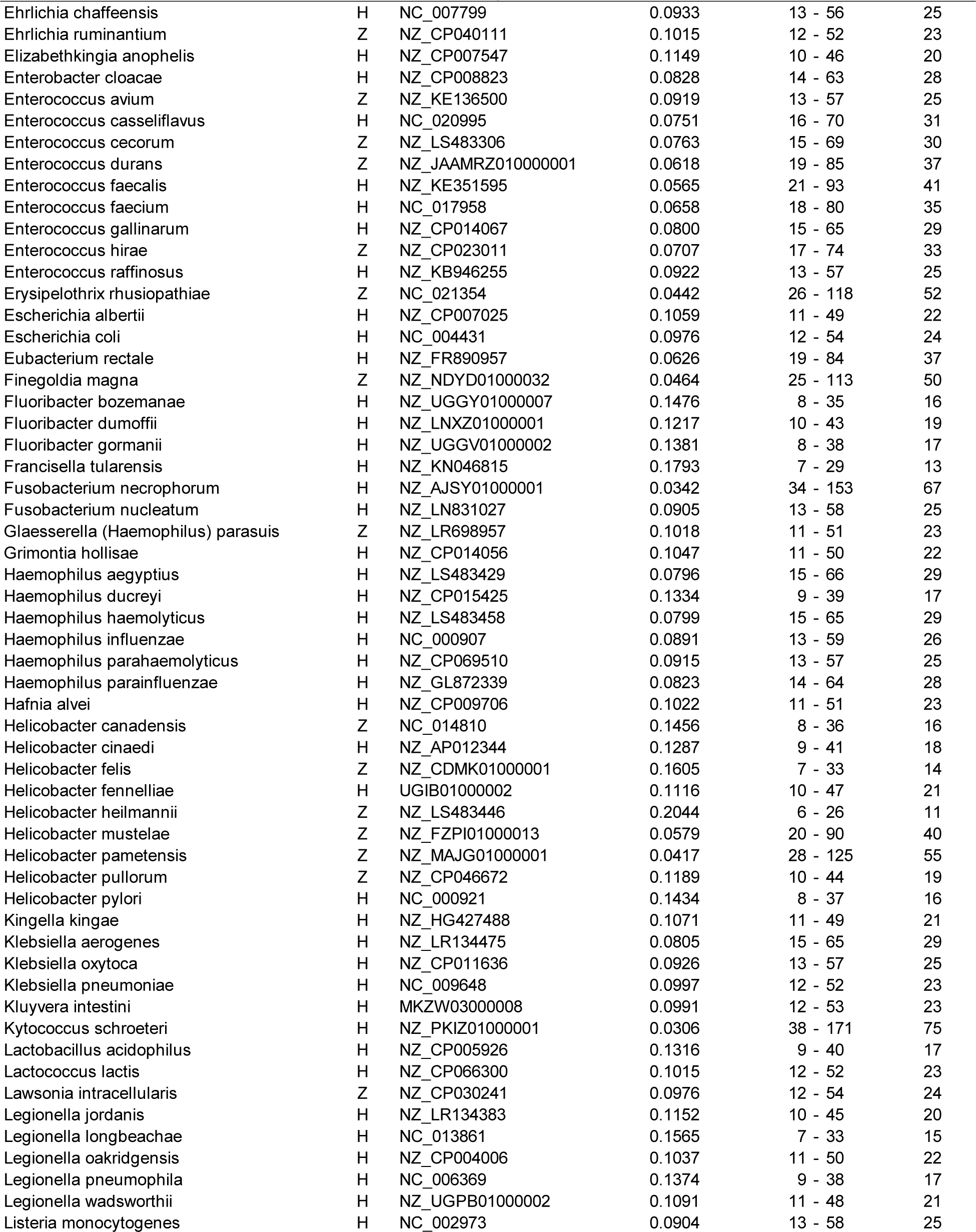
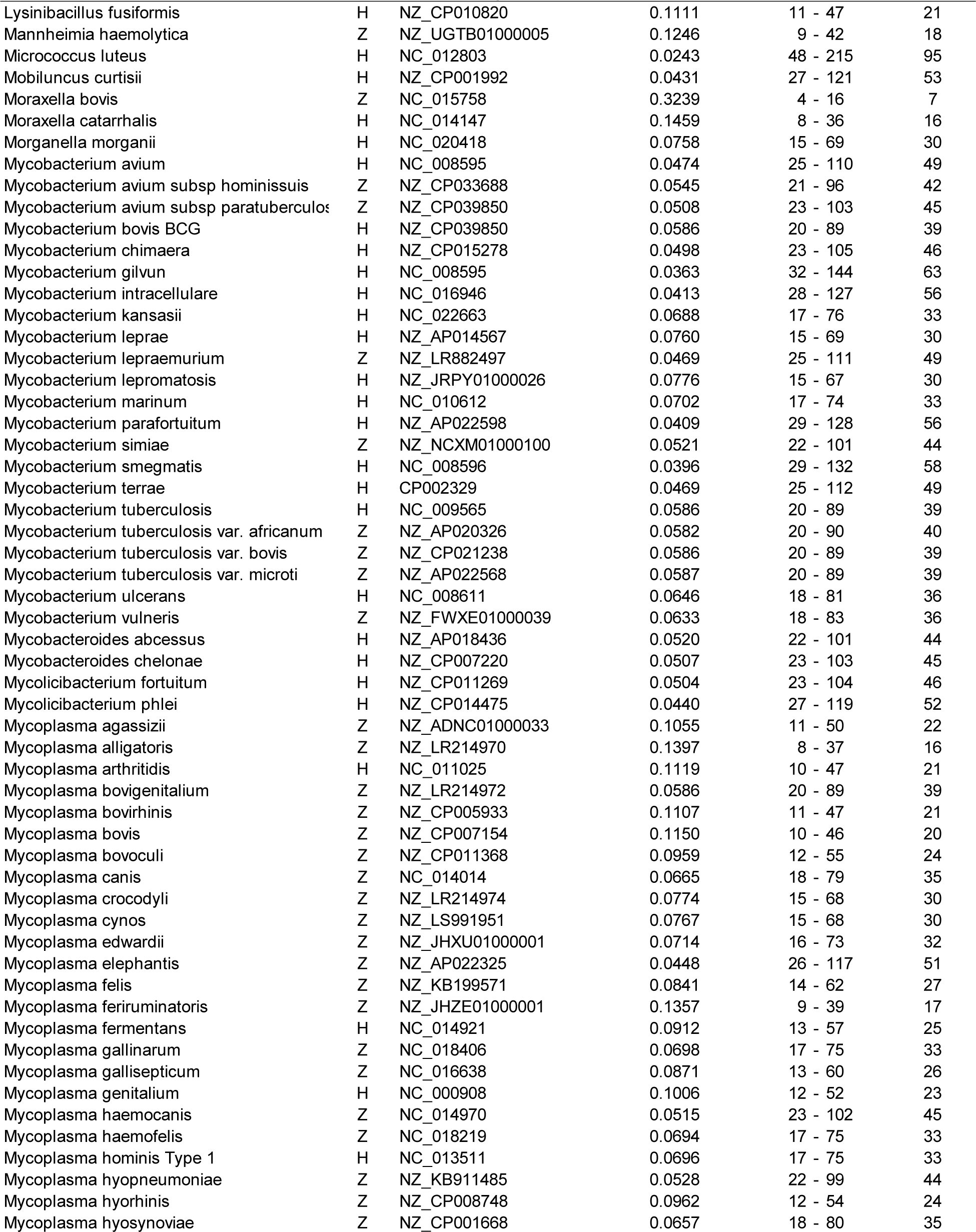
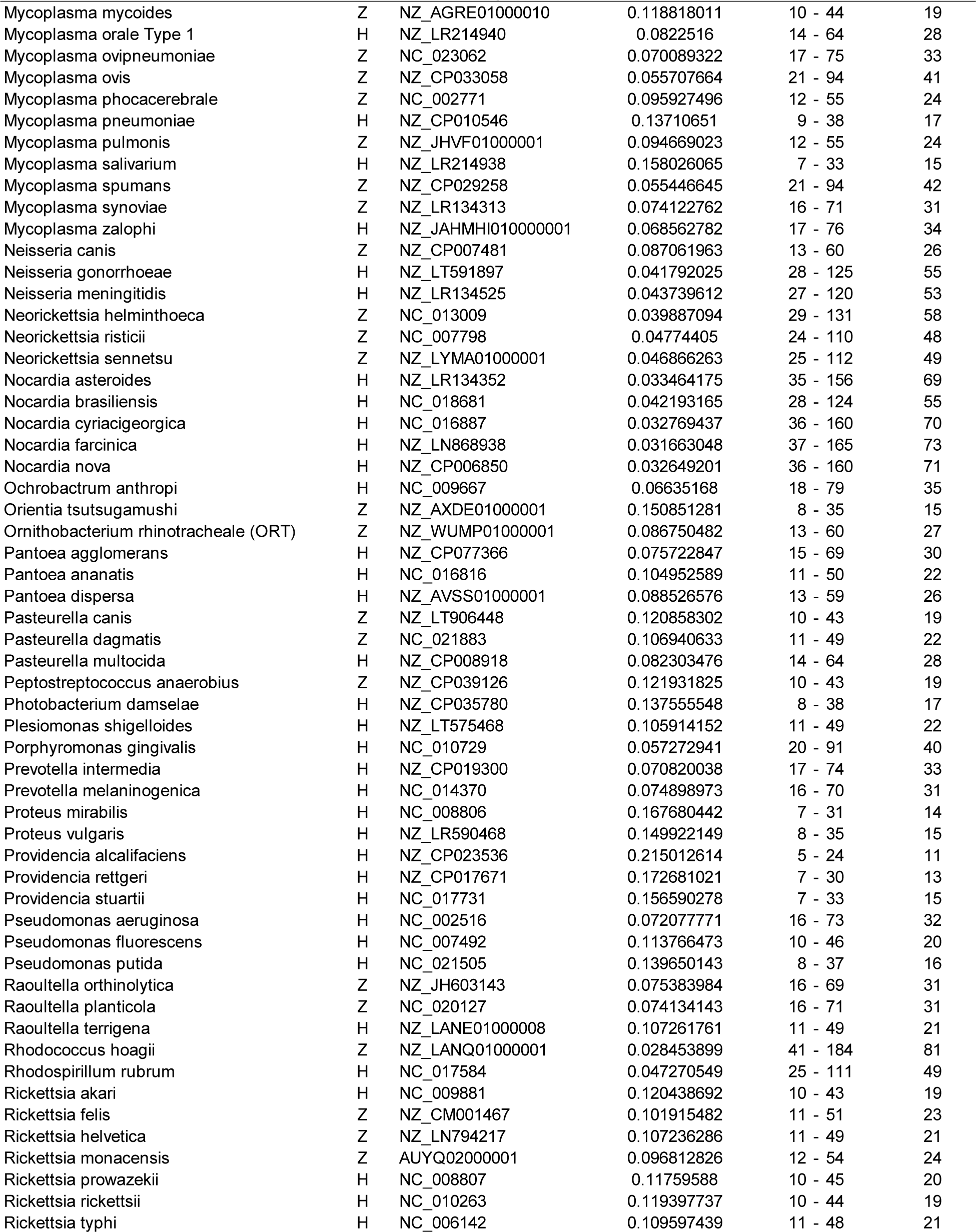
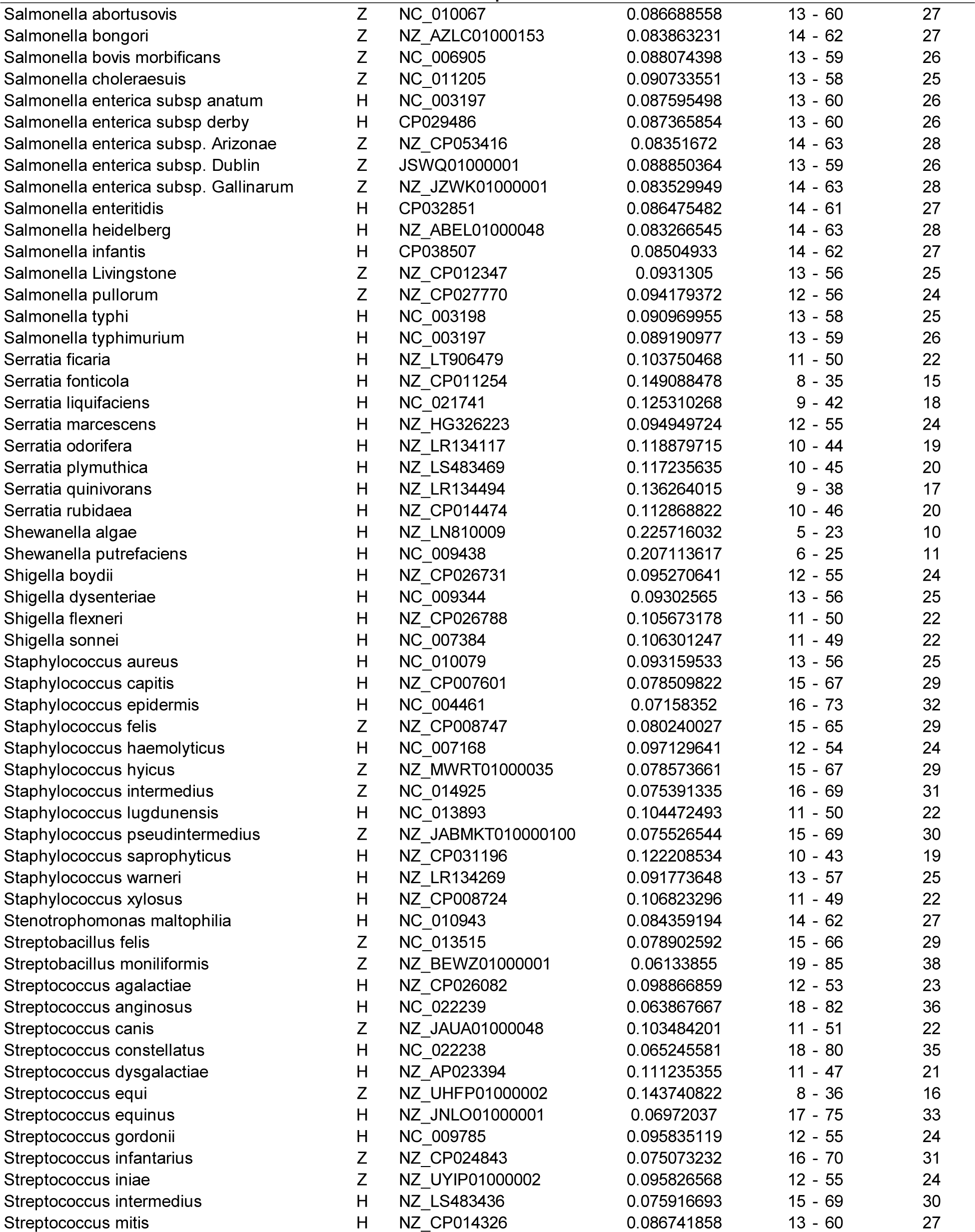
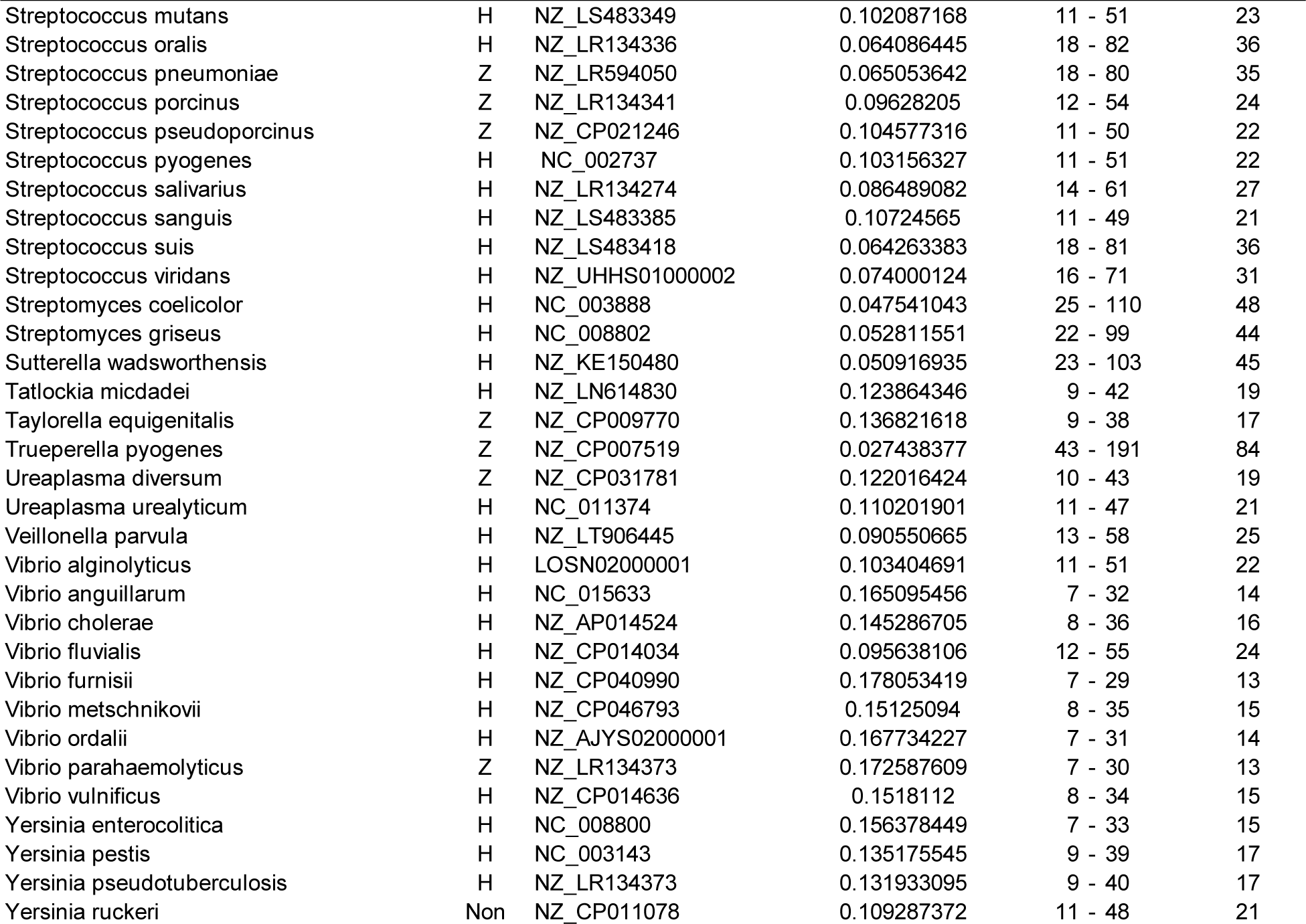
Bacterial Pathogen Ultraviolet Susceptibility Predictions.

The genomic parameters used in the model were further studies to evaluate their relative contribution to the model. The contributions were analyzed as stand-alone correlations and the results are shown in Figure 3. Only the Hot Spots were used for this model and no correlations can be made for the individual dimers. The results indicate that the combined Hot Spots (TTh+TCh+CTh+CCh+YRh) were the single strongest predictor with a 70% Pearson correlation coefficient.

**Figure 3:**
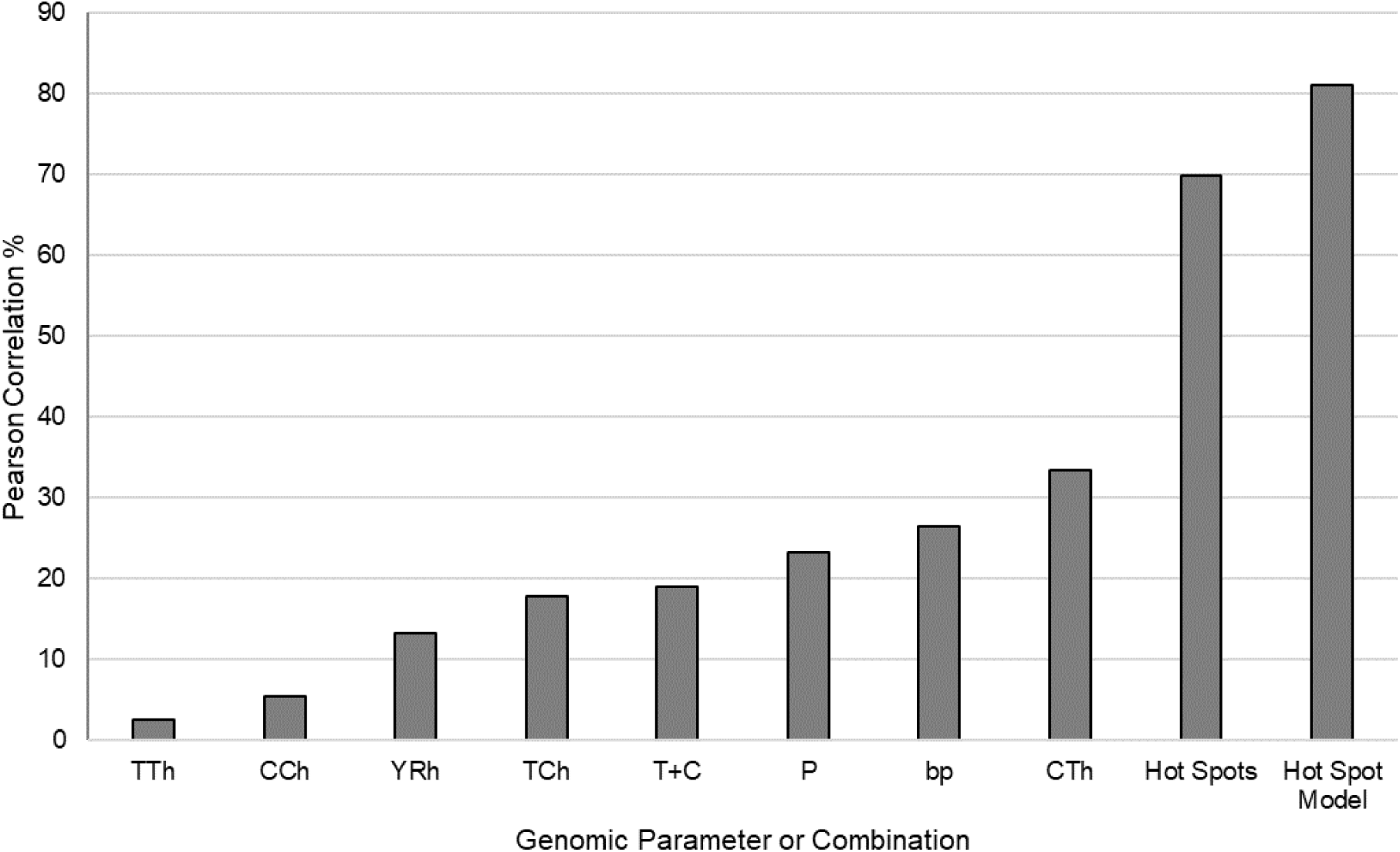
Pearson Correlation coefficients for genomic parameters in isolation and in combinations of parameters. Hot Spots are the sum of all hot spots (TTh+TCh+CTh+CCh+YRh).

A graph of the Hot Spots was made and the results are presented as a circular genome for the bacteria *Borrelia afzeli* in Figure 4. In this graph only the highest hyperprimer values for the first 4000 bp are shown for simplicity. Figure 4 shows the pyrimidine hot spots in red and the purine hot spots in yellow. It is worth noting that of the hundreds of indicated Hot Spots only a few would be necessary to inactivate the pathogen. The purine hot spots are significant because photoreactivation enzymes may repair pyrimidine dimers but they cannot repair purine dimers. Purine dimers may therefore be more fatal to the pathogen than pyrimidine dimers.

**Figure 4:**
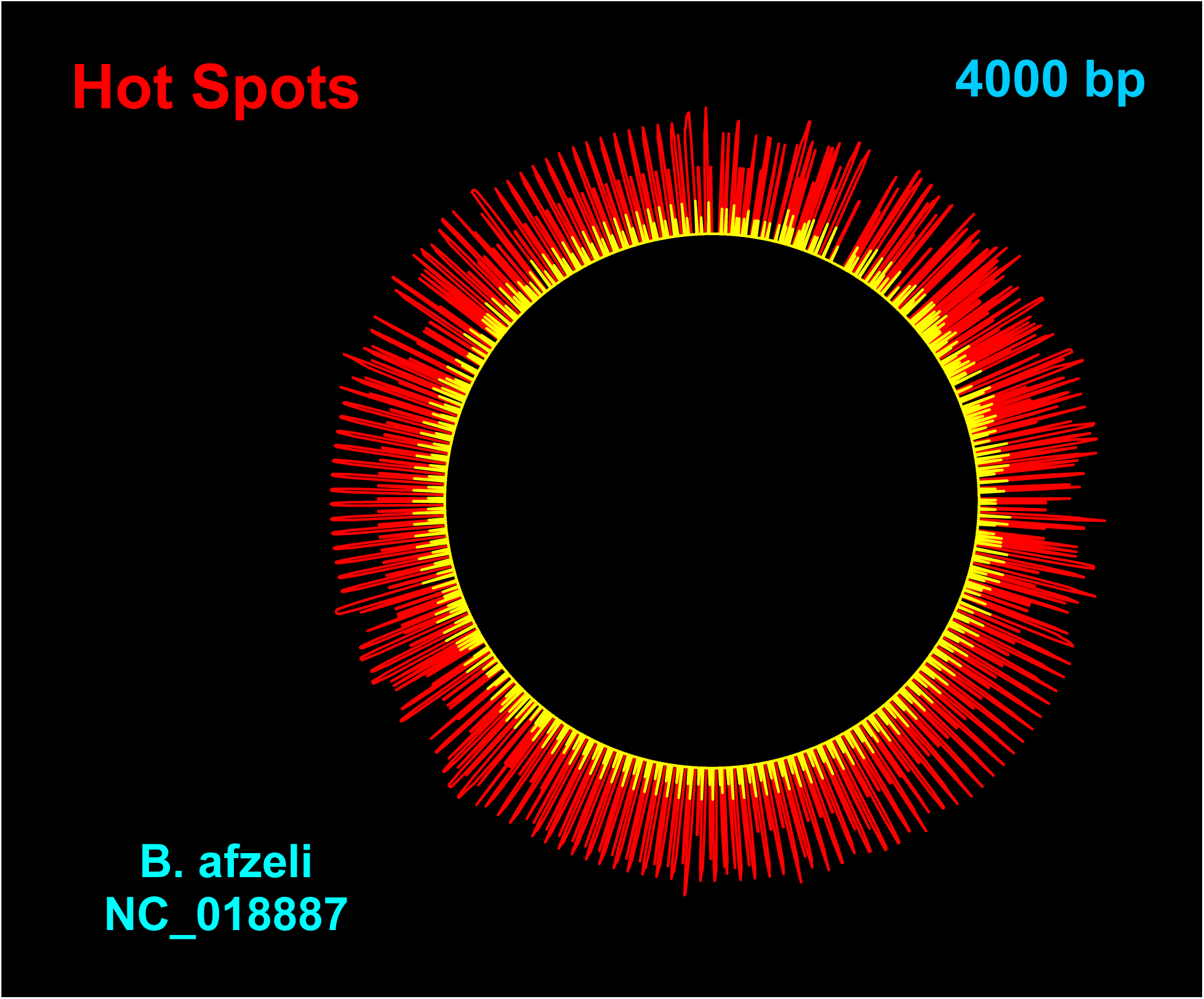
Example of hot spots highlighted on the circular genome of *Borrelia afzeli*. Red signifies pyrimidine hot spots where three or more pyrimidines are clustered. Yellow signifies hot spots at purine bases.

## Discussion

The present genomic model improves upon the results presented previously by the author (Kowalski 2009, 2009a, 2009b, 2009c). The current Hot Spot bacteria model incorporates many more studies than previous models and more scrutiny was placed on calculating the UV rate constants but the results are not drastically different from previous attempts. While there are clearly a great many variations possible in the modeling approach and in the selection and adjustment of parameters, the final predictions of UV rate constants have changed little, suggesting they limited sensitivity to the approach.

Other attempts to develop genomic models for viruses have been published and these have employed much the same genomic motifs as specified in the current model and have also produced similar predictions of UV rate constants (or D90 values) as the author’s virus genomic models (Rockey 2021, Pendyala 2020). Although a fundamentals-based genomic model has been elusive, it appears that empirically based genomic models are adequate to generate predictions of UV sensitivity for all types of pathogens. Genomic modeling has also been used by the author to successfully predict the UV sensitivity of individual antibiotic resistant genes (Destiani 2017).

We invite researchers to challenge these predictions, especially those bacteria that have no prior studies, and also for the four outlying bacteria. The predicted values for UV rate constants can act as a target for the experimental design and should enable researchers to seek more accurate results. Because of the fact that the UV rate constant must be absolutely defined by the species genome, the predictions provided here may act as a standard by which to gauge results from future experiments.

It was clear from reviewing all studies that there was a lack of consistency in the test methods and this surely resulted in scattering of the results and reduced predictive performance of the model. Problems were observed in many studies related to how the UV dose was established. Researchers should take care to use photosensors properly. In general, if a collimated beam apparatus is not used, the distance from the photosensor to the lamp should be on the order of one lamp length. In addition, many studies used low levels of UV irradiance and this produced a shoulder in the survival curve. The shoulder will distort any D90 value estimated from the data. It is suggested that researchers use irradiance levels of at least 1 W/m^2^ (100 µW/cm^2^) to avoid shoulders in the results and allow more accurate determination of the first stage rate constant.

A standardized test for UV rate constant evaluation is needed. Genomic modeling makes it possible to compute UV rate constants to six decimal places for bacteria, which have genomes on the order of 1M bp. A standard test capable of resolving UV rate constants to six decimal places, which represents six logs of reduction, may require the use of bacteria monolayers to avoid tailing in the survival curve, which reduces measurement accuracy.

The current method makes it possible to quickly evaluate the UV susceptibility of any emerging pathogen once the genome is published. The calculations could conceivably be incorporated into the NCBI Genome database to provide instant results once a genome is sequenced. Future research will address improved genomic modeling of viruses and fungi and investigation will be conducted into whether the Hot Spot model may have applications in predicting susceptibility to skin cancer.

